# Single-cell multi-omics analysis reveals cooperative transcription factors for gene regulation in oligodendrocytes

**DOI:** 10.1101/2024.06.19.599799

**Authors:** Jerome J. Choi, John Svaren, Daifeng Wang

## Abstract

Oligodendrocytes are the myelinating cells within the central nervous system. Many oligodendrocyte genes have been associated with brain disorders. However, how transcription factors (TFs) cooperate for gene regulation in oligodendrocytes remains largely uncharacterized. To address this, we integrated scRNA-seq and scATAC-seq data to identify the cooperative TFs that co-regulate the target gene (TG) expression in oligodendrocytes. First, we identified co-binding TF pairs whose binding sites overlapped in oligodendrocyte-specific regulatory regions. Second, we trained a deep learning model to predict the expression level of each TG using the expression levels of co-binding TFs. Third, using the trained models, we computed the TF importance and TF-TF interaction scores for predicting TG expression by the Shapley interaction scores. We found that the co-binding TF pairs involving known important TF pairs for oligodendrocyte differentiation, such as SOX10-TCF12, SOX10-MYRF, and SOX10-OLIG2, exhibited significantly higher Shapley scores than others (t-test, p-value < 1e-4). Furthermore, we identified 153 oligodendrocyte-associated eQTLs that reside in oligodendrocyte-specific enhancers or promoters where their eGenes (TGs) are regulated by cooperative TFs, suggesting potential novel regulatory roles from genetic variants. We also experimentally validated some identified TF pairs such as SOX10-OLIG2 and SOX10-NKX2.2 by co-enrichment analysis, using ChIP-seq data from rat peripheral nerve.

## Introduction

Oligodendrocytes play key functional roles in the central nervous system (CNS) function, including that they are responsible for myelination^1,2^. Myelination is a complex neurodevelopmental process that begins during brain development in the third trimester of pregnancy and increases steadily during childhood, but it can also be dynamically regulated in the context of learning and diseases affecting the mature CNS^3,4^. Also, Oligodendrocyte dysfunction and myelin abnormalities have been reported in CNS disorders^2,5,6^. Multidirectional interactions between neuronal and glial cells are required for CNS function^7^, including interactions between oligodendrocytes and neurons through myelination^8^. Therefore, it is critical to better understand the functions and roles of oligodendrocytes and myelin.

Gene expression of oligodendrocyte development from oligodendrocyte progenitor cells (OPC) is governed by complex gene regulatory mechanisms involving transcription factors (TFs)^3,4^. TFs often work in a combinatorial fashion to regulate gene expression from regulatory elements^9,10^. For example, some TFs such as SOX10 and OLIG2 cooperate during the induction of genes for differentiation and myelin formation^11–14^. Enhancers potentially enhance transcription levels from promoters and transcriptional start sites (TSS), and much of the regulatory code that drives cell type-specific gene expression resides in distal regulatory elements, enhancers. Especially, some active enhancers are associated with the gene expression that characterizes cell identity and functions^15^. Thus, it is important to identify active oligodendrocyte-specific enhancers and the co-binding TFs that are responsible for their activity.

Next-generation sequencing technologies, including single-cell RNA sequencing (scRNA-seq) and the assay for transposase-accessible chromatin sequencing (scATAC-seq), have provided important insights into cell-type-specific gene regulation. Recent functional genomic resources such as PsychENCODE^16^ and GTEx^17^, and emerging tools for integrating multi-omics data enable creating cell-type-level gene regulatory networks (GRNs) linking TFs and their binding sites (TFBS), regulatory elements to target genes (TGs). Those networks can reveal the regulatory roles of cell-type TFs via regulatory elements. Moreover, additional bioinformatic tools such as SCENIC^18^, Signac^19^, and scGRNom^20^ predict cell type gene regulatory networks to explain potential TF-TG relationships. However, most of these studies and tools focus on relationships between individual TFs and TGs instead of TF-TF interactions and their effects on TG expression. Consequently, due to the lack of tools, the mechanistic roles of cooperative TFs in establishing cell type-specific gene regulation remain uncharacterized.

## Results

### Deep learning and single-cell multiomics for identifying cooperative transcription factors in oligodendrocytes

In order to predict cooperative TFs involved in oligodendrocyte gene regulation, we designed an analytical framework to analyze single-cell multi-omics data for a single cell type from human brain single cell data (**Fig.1**, Methods and Materials). Briefly, we first used scATAC-seq data with peak-to-gene links^21^. Second, among the regulatory regions for different cell types, we filtered those for oligodendrocytes and identified transcription factor binding sites (TFBSs) and co-binding transcription factor (TF) pairs from motif co-occurrence and co-enrichment analyses. Third, we trained deep neural networks (DNNs) to predict the expression levels of the TGs and measure interaction effects between co-binding TFs on the expression levels of TGs using gene expression from scRNA-seq data^22^ and computed SI scores for co-binding TF pairs and found cooperative TF pairs. Fourth, we built a gene regulatory network based on SI scores for co-binding TF pairs. Lastly, as an independent validation, we mapped oligodendrocyte eQTLs onto the regulatory regions where cooperative TF pairs exist. For another independent validation, we converted the regulatory regions where cooperative TF pairs exist to the rat genome assembly, and measured the density of ChIP-seq signal and implemented a co-enrichment analysis.

**Fig. 1:**
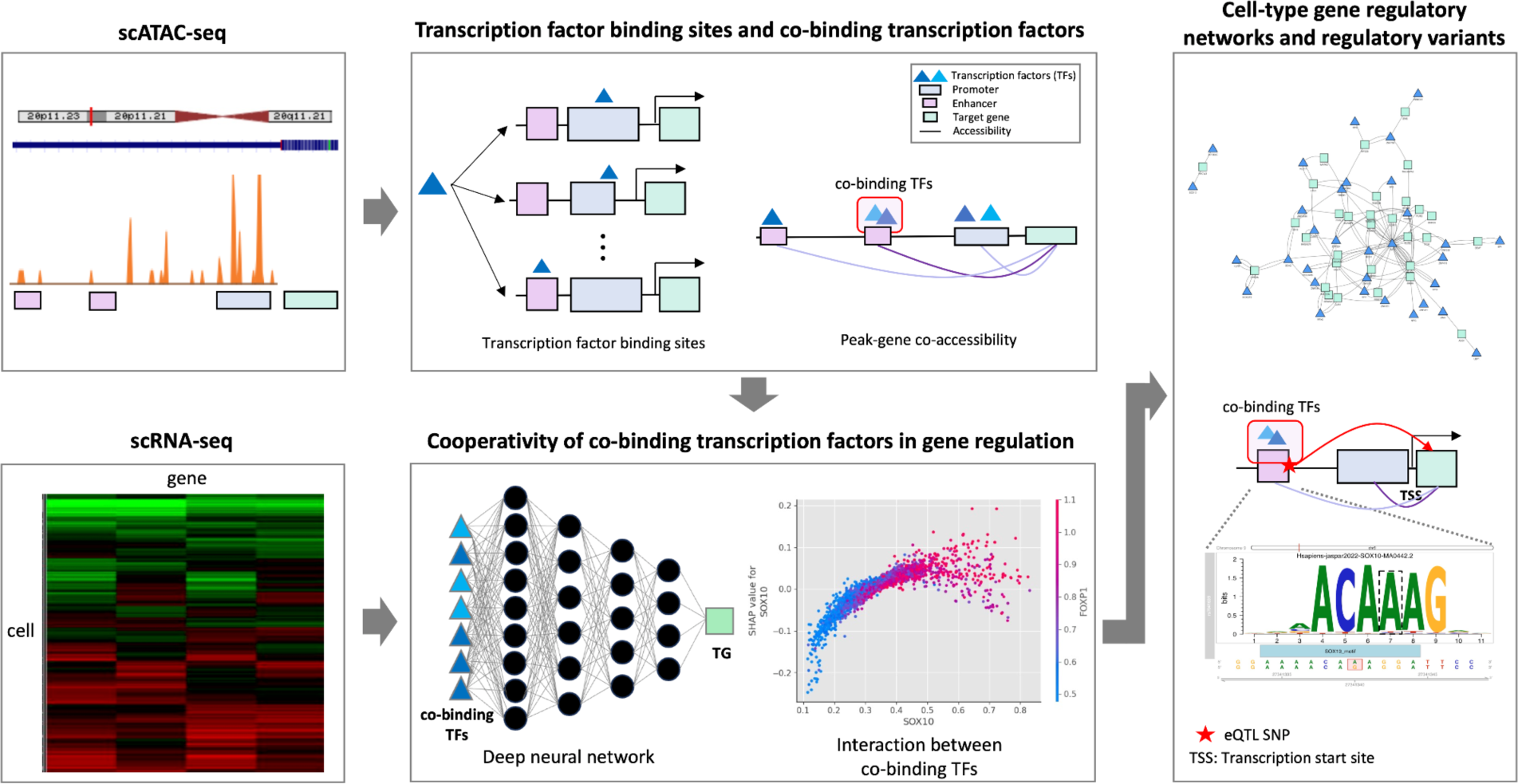
Deep learning and single-cell multiomics for identifying cooperative transcription factors in oligodendrocytes. Inputs for the pipeline are scATAC-seq peak-gene links and scRNA-seq. It infers transcription factor binding sites (TFBSs) in regulatory regions and identifies co-binding TF pairs. Then, it measures cooperativity of co-binding TFs by predicting TF-TG relationships for the levels of expression using deep learning models and Shapley interaction scores. It outputs a gene regulatory network linking co-binding TF pairs with their TGs and regulatory variants on the regulatory regions where co-TFs have their binding sites.

### Identification of the co-binding transcription factors in oligodendrocyte-specific regulatory regions

First, we identified a set of 787 oligodendrocyte-differentially accessible and oligodendrocyte-specific regulatory regions by comparison of oligodendrocyte scATAC-seq data to other brain cell types. In this set, we identified 958 motifs for inferred TFBSs using the JASPAR database. Second, we implemented co-occurrence analysis and co-enrichment analysis for 458,403 possible TF pairs in the regulatory regions. We removed TF pairs from the same families and applied a cutoff (<0.1) for false discovery rate (FDR) yielding 8,101 co-binding TF pairs. There were 206 TFs that have co-binding TFs linked to 445 TGs (Supplementary Data) that are oligodendrocyte specific in 643 regulatory regions (**Fig. 2a**). We annotated the regulatory regions to categorize them into promoters (32.5%) and enhancers (67.5%) (**Fig. 2b)**.

**Fig. 2:**
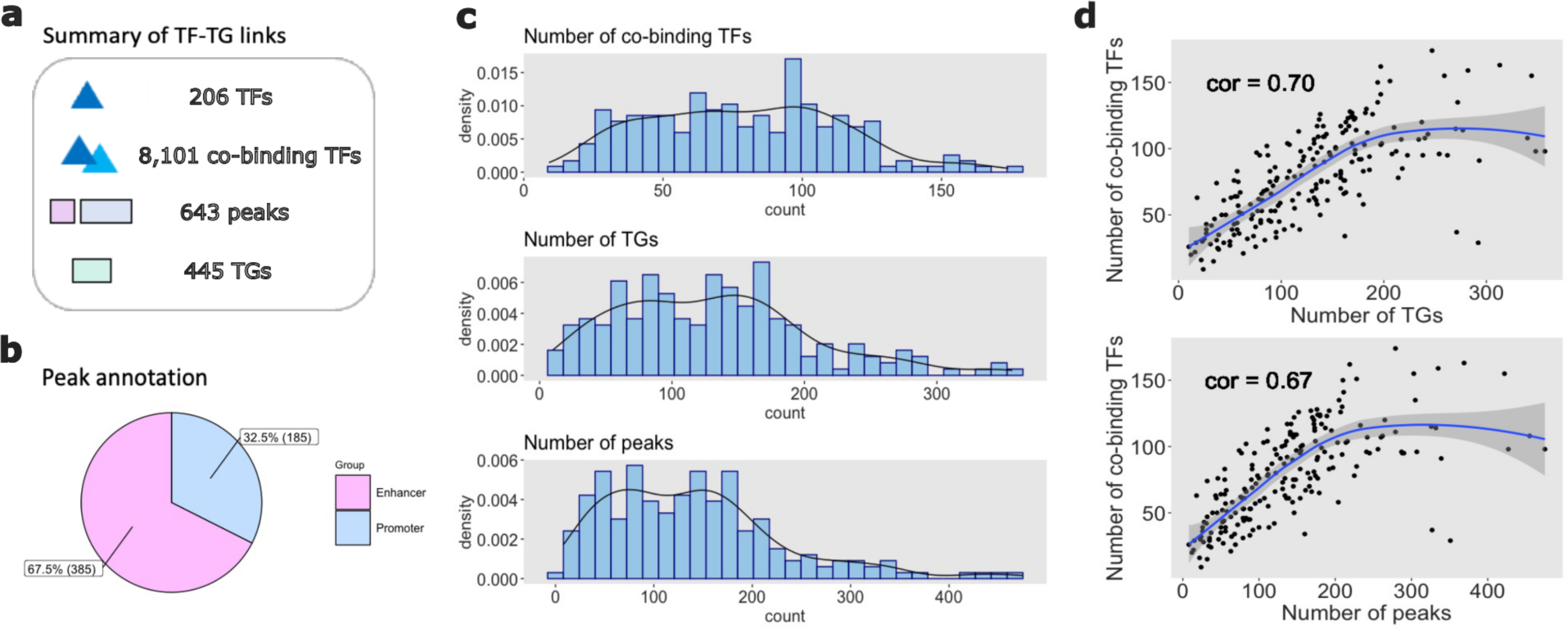
Distribution and correlation of numbers of co-binding transcription factors, target genes, and peaks for individual transcription factors, peak annotation, and summary statistics for transcription factor-target gene links. **a** Summary statistics of transcription factor (TF)-target gene(TG) links. **b** Peak annotation. **c** Distributions of number of co-binding TF), TGs, and peaks for individual transcription factors. **d** Correlations between the numbers of co-binding transcription factors and target genes and the numbers of co-binding TFs and the number of peaks.

The density plots show the distributions of the number of co-binding TF pairs, the number of TGs, and the number of peaks for individual TFs (**Fig. 2c**) that are co-bound to other TFs. Most of the TFs have 50 to 103 co-binding TFs (median = 78). The distribution of the number of TGs for TFs is right-skewed, and many TFs have 76 to 172 TGs linked. The distribution of the number of peaks for TFs is also right-skewed, and the most frequent intervals were between 75 to 180 peaks. Additionally, other density plots show the distributions of the number of TGs and the number of peaks for co-binding TF pairs (**Fig. S1**) and bar plots displayed the numbers of co-binding TFs, TGs, and peaks for individual TFs by their family categories (**Fig. S2**). Co-binding TFs have 4 to 115 TGs (median = 59) and 4 to 123 peaks (median = 56) and the most frequent motifs are associated with TF families with C2H2 zinc finger, bZip, and bHLH DNA-binding domains.

We computed Pearson correlation coefficient (*r*) to measure correlations between the number of co-binding TFs and the number of TGs and the number of co-binding TFs and the number of peaks for individual TFs (**Fig. 2d**). The number of co-binding TFs and the number of TGs for individual TFs are strongly positively correlated (*r* = 0.70). It suggests that TFs that are co-bound to other TFs tend to have more TGs linked to them. The number of co-binding TFs and the number of peaks for individual TFs are also strongly positively correlated (*r* = 0.67). It shows that co-binding TFs may exist in many different peaks.

### Oligodendrocyte gene expression relationships between transcription factors and target genes

A single cell study identified the unique gene expression profile of oligodendrocytes compared to other brain cell types^22^, as shown by the two dimensional Uniform Manifold Approximation and Projection (UMAP) space after computing latent representations of the neighborhood graph (**Fig. 3a**). In order to focus on oligodendrocyte-specific mechanisms of gene regulation, we conducted differential expression testing using 17,946 genes and 20,191 metacells and identified 4,387 differentially expressed genes (DEGs) for oligodendrocytes. We found 445 TGs out of 507 TGs of oligodendrocyte-specific regulatory elements (88%) were DEGs for oligodendrocytes. Subsequently, we conducted enrichment analysis for these 445 TGs revealing their involvement in crucial biological processes for oligodendrocytes, such as oligodendrocyte development, oligodendrocyte differentiation, and myelination (**Fig. 3b**).

**Fig. 3:**
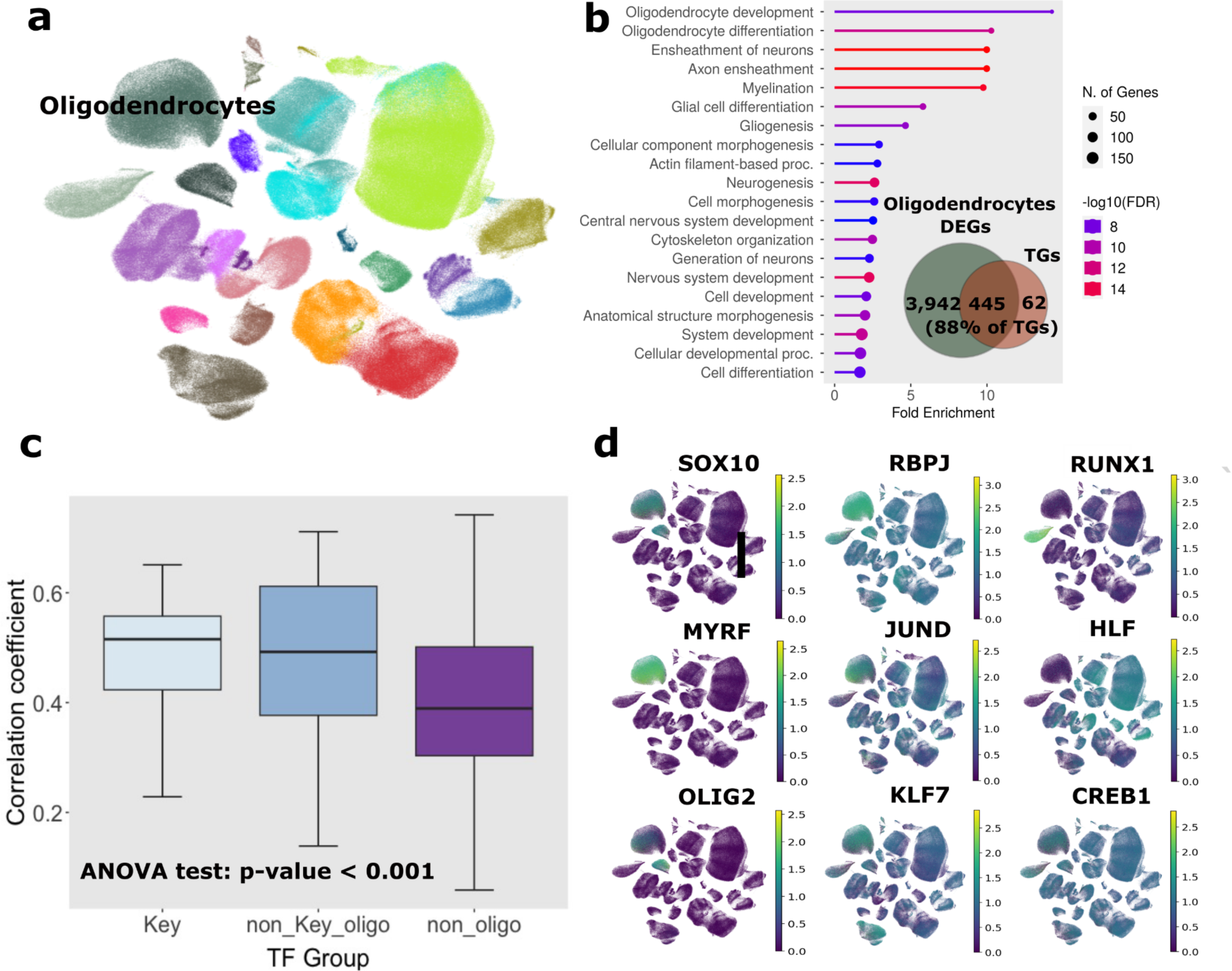
Oligodendrocyte gene expression relationships between transcription factors and target genes. **a** UMAP for eighteen cell type in middle temporal gyrus region, **b** Enrichments analysis for target genes that are oligodendrocyte-specific, **d** Expression level comparison between three categories of transcription factor pairs: oligodendrocyte key transcription factor pairs, non-key oligodendrocyte-specific transcription factor pairs, and non-oligodendrocyte-specific transcription factor pairs, and **d** UMAPs for examples of the three categories in **c** (each column is an example for each category).

We categorized TFs into oligodendrocyte key TFs, oligodendrocyte-specific non-key TFs, and non-oligodendrocyte-specific TFs using oligodendrocyte DEGs and the list of key TFs. ‘Oligodendrocyte-specific key TFs’ are oligodendrocyte DEGs and key TFs, ‘oligodendrocyte-specific non-key TFs’ are oligodendrocyte DEGs but not key TFs, and ‘non-oligodendrocyte-specific TFs’ are neither oligodendrocyte DEGs nor key TFs. The key oligodendrocyte TF’s were defined based on mouse loss-of-function studies that have shown that specific TF’s are critical for oligodendrocyte differentiation. The key TF’s include SOX10^23^, SOX2,^24,25^ SOX8,^26^ MYRF^27^, OLIG1^28^, OLIG2^29^, TCF7L2,^30,31^ ZNF24^32^, NKX2.2^33^, and NKX6.2^34^.

Each of 206 TF, who have co-binding TFs, regulates a different set of TGs, and we computed correlations between each TF in the three categories and its TGs. An ANOVA test shows that there is a significant difference in means among the three categories (p-value < 0.001) (**Fig. 3c**). We generated UMAPs, selecting three TFs as examples for each category. Oligodendrocyte-specific key TFs such as SOX10, MYRF, and OLIG2 are specifically expressed in oligodendrocytes. Oligodendrocyte-enriched non-key TFs, including RBPJ, JUND, and KLF7, are expressed in multiple cell types but are more highly expressed in oligodendrocytes. Non-oligodendrocyte-specific TFs, such as RUNX1, HLF, and CREB1, are not specifically expressed in oligodendrocytes (**Fig. 3d**).

### Deep learning and Shapley interaction scores to measure cooperativity of co-binding transcription factors

To understand the complex relationships between TFs for predicting TGs, we built deep learning models. We trained a deep learning model for each of the 445 TG. Each model used the expression levels for the 202 TFs that have co-binding TFs to predict a TG expression level. We used seven hidden layers in each DNN (**Fig. 4a**). We calculated the mean and standard deviations of the interaction scores for all TGs for each co-binding TF pair and for co-binding TF pairs that have high variations (coefficient of variance > 0.5). Using a trained model and a hold-out test dataset, we computed SI scores for TFs in each DNN. Additionally, we determined the percentile SI score for all co-binding TF pairs. Then, a t-test to compare the mean values for the percentile SI scores of key co-binding TF pairs and non-key co-binding TF pairs revealed a significant difference between the two groups (p-value < 0.0001) (**Fig. 4b**).

**Fig. 4:**
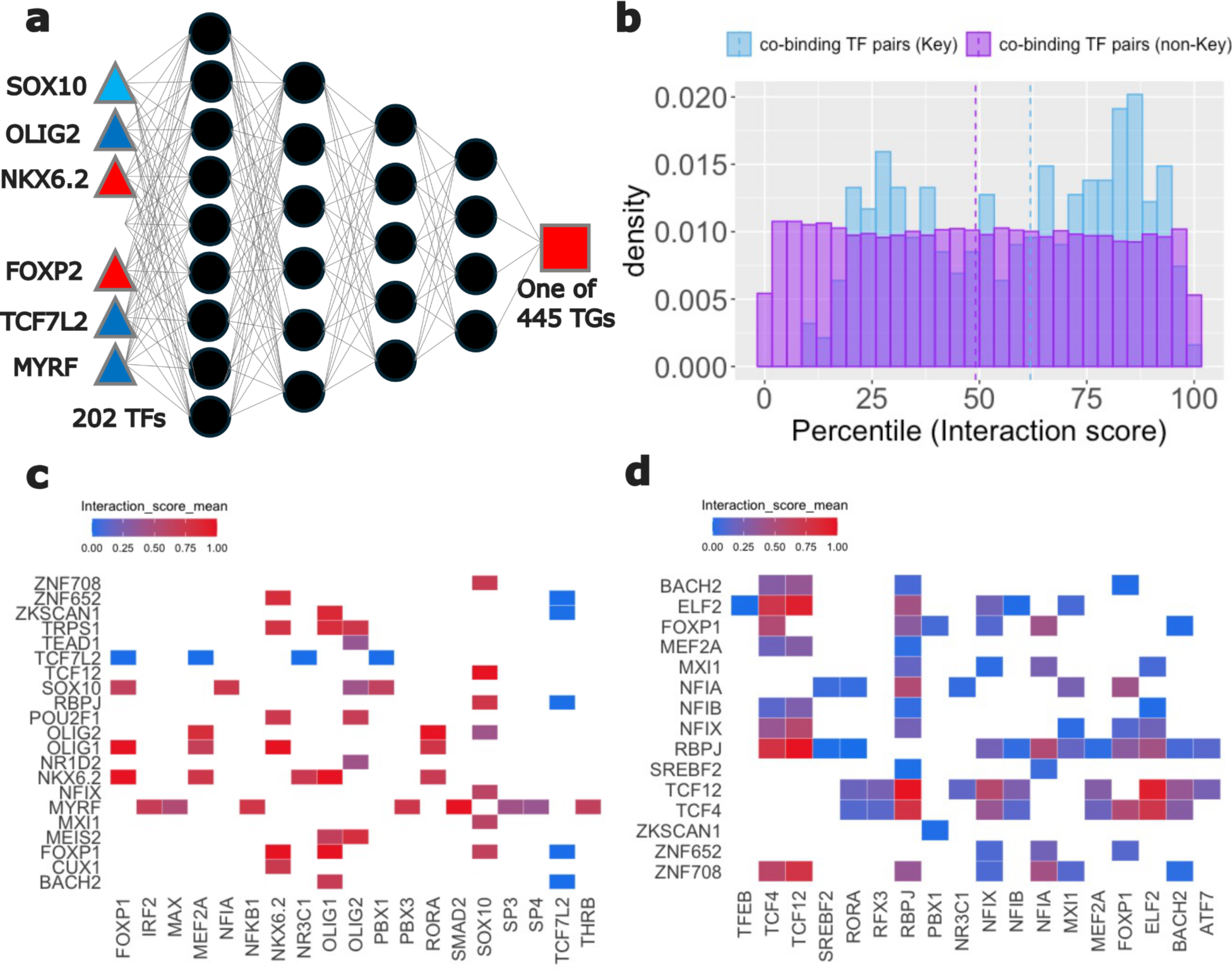
Cooperative transcription factor pairs by Shapley interaction scores. **a** Deep learning architecture, **b** T-test for the interaction score distributions of key-transcription factor pairs and non-key transcription factor pairs, **c** Top fifty-five interaction scores for key transcription factor pairs, and **d** Top fifty-five interaction scores for non-key transcription factor pairs.

We selected the top nine interacting pairs for each key co-binding TF pair, such as SOX10, MYRF, OLIG1, OLIG2, NKX6.2, and TCF7L2, and generated a heatmap for their SI scores scaled from 0 to 1 (**Fig. 4c**). Similarly, we chose the top fifty-four interacting co-binding TF pairs for non-key TFs and created another heatmap for their SI scores scaled from 0 to 1 (**Fig. 4d**). We noticed that the SI scores for key-TF co-binding pairs have higher values than those for non-key co-binding TF pairs.

We also validated our model prediction performance for one TG, *MBP*, using additional data^35^ (**Fig. S3**). We regressed the scaled actual values on the scaled predicted values. For our primary dataset, we obtained an R-squared of 0.81 and a *r* of 0.90 (**Fig. S3a**). Furthermore, when analyzing another dataset, we observed an R-squared of 0.69 and a *r* of 0.83, affirming the predictive capability of our model architecture (**Fig. S3b**).

### Oligodendrocyte gene regulatory network analysis for cooperative TF pairs and transcription factor hierarchy

We chose one pair of co-binding TF with the highest interaction scores from six key co-binding TF pairs, including SOX10, MYTF, OLIG1, OLIG2, NKX6.2, and TCF7L2. We built a gene regulatory network (GRN) for these cooperative TF pairs and their TGs that are co-regulated by them (**Fig. 5a**). We found that a TG, *CALD1*, is co-regulated by three key cooperative TF pairs, SOX10-TCF12, RORA-OLIG2, and FOXP1-NKX6.2 and another TG, *PPP1R16B*, is co-regulated by three key cooperative TF pairs, RORA-OLIG2, FOXP1-NKX6.2, and FOXP1- OLIG1. There are other TGs in the GRN that are co-regulated by two pairs of cooperative TFs.

**Fig. 5:**
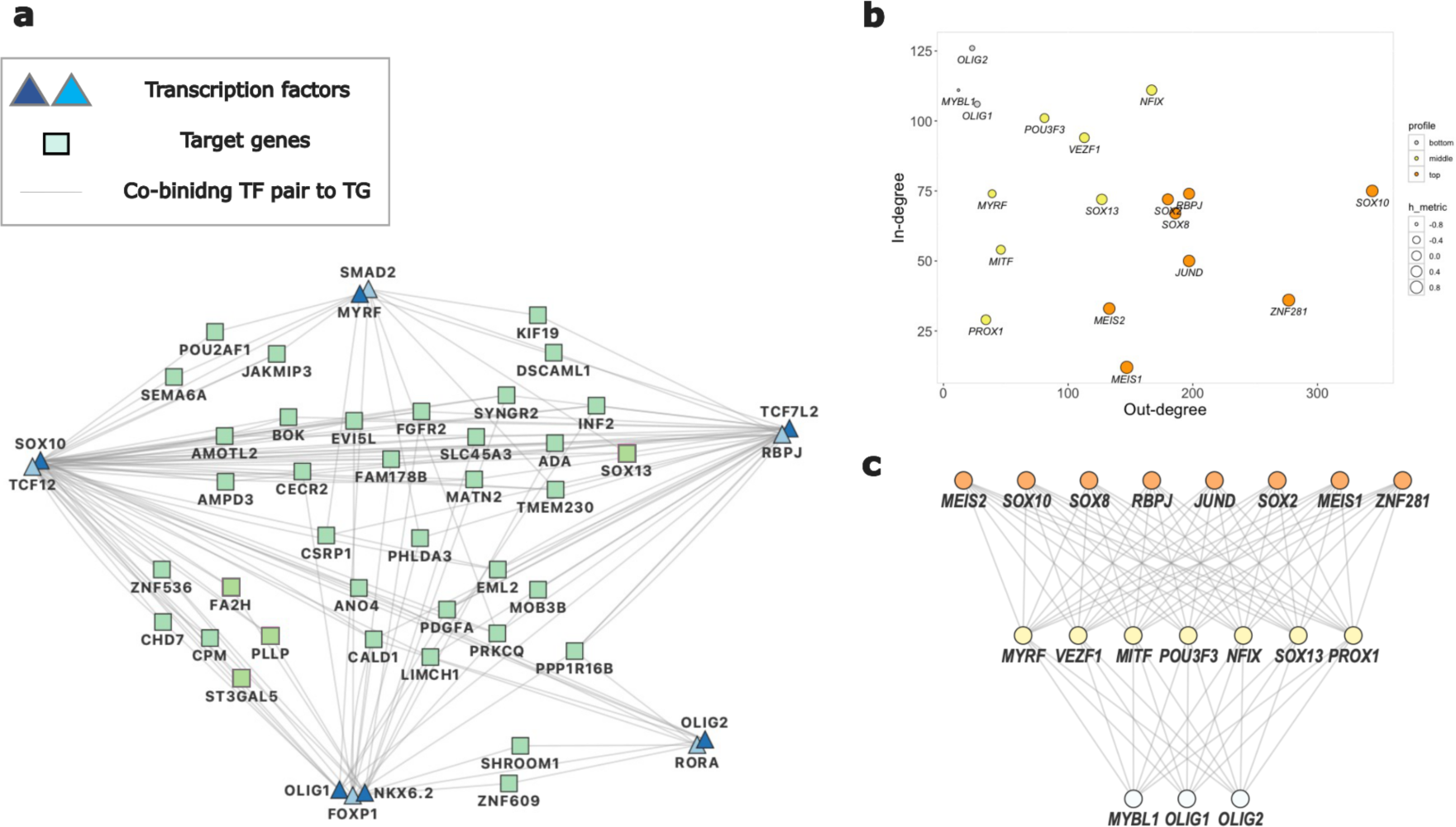
Gene regulatory network and transcription factor hierarchy. **a** Gene regulatory network for six key cooperative transcription factor pairs with the highest interaction scores, **b** A plot of in-degree (I) vs out-degree (O) for the transcription factors that have I and O in the gene regulatory network, and **b** Transcription factor hierarchy. Each node depicts a transcription factor. In **b** and **c**, the edges colored in orange are the top-level (master) regulators, the edges colored in orange are the middle-level regulators, and the edges colored in yellow are the bottom-level regulators.

We computed in-degree and out-degree for eighteen TFs that can also be TGs at the same time since TF feed forward and feedback loops are common (**Fig. 5b**). Then, we conducted a TF hierarchy analysis and found eight top-level regulators (Fig. 5c), called ‘Master regulators’, including SOX10, SOX2, and SOX8, which are key TFs that are known to play critical roles in oligodendrocyte differentiation.^24–26^ The other five master regulators, MEIS1, MEIS2, RBPJ, JUND, and ZNF281, are categorized as oligodendrocyte-specific non-key TFs in **Fig. 3c**. All TFs that are middle-level regulators and bottom-level regulators, except for MYRF, are categorized as oligodendrocyte-specific non-key TFs. MYRF is one of the key TFs which is specifically activated in myelinating oligodendrocytes. PROX1 has been identified as being important for oligodendrocyte differentiation^36,37^. Most of these eighteen TFs are expressed in both oligodendrocytes and OPCs (**Fig. S5**). It provides evidence that oligodendrocyte differentiation is pre-set in OPCs^38^.

### Independent validation for cooperative TFs

**eQTL mapping.** As an independent assessment of the regulatory regions, we mapped oligodendrocyte eQTLs^39^ onto oligodendrocyte-specific regulatory regions to explain the causal relationships between the expression levels of the co-binding TF pairs we identified and their target genes (TGs). Notably, it provides evidence of causation if the eQTL genes and TGs are identical where co-binding TF pairs occur, indicating that these co-binding TF pairs are co-regulating TG expressions.

First, among 4.8 million oligodendrocytes eQTLs, we filtered 2 million significant (FDR < 0.05) eQTLs. Second, we mapped these significant eQTL SNPs (eSNPs) onto oligodendrocyte-specific regulatory regions (**Fig. 6a**). In total, 384 eSNPs and 160 eQTL genes (eGenes) were mapped onto 188 regulatory regions. Among these, 373 eSNPs and 153 eGens (TGs) were found in 179 regulatory regions associated with key TF pairs. Enrichment analysis for TGs indicates their strong involvement in biological processes such as oligodendrocyte development, myelination, and oligodendrocyte differentiation. (**Fig. 6a**)

**Fig. 6:**
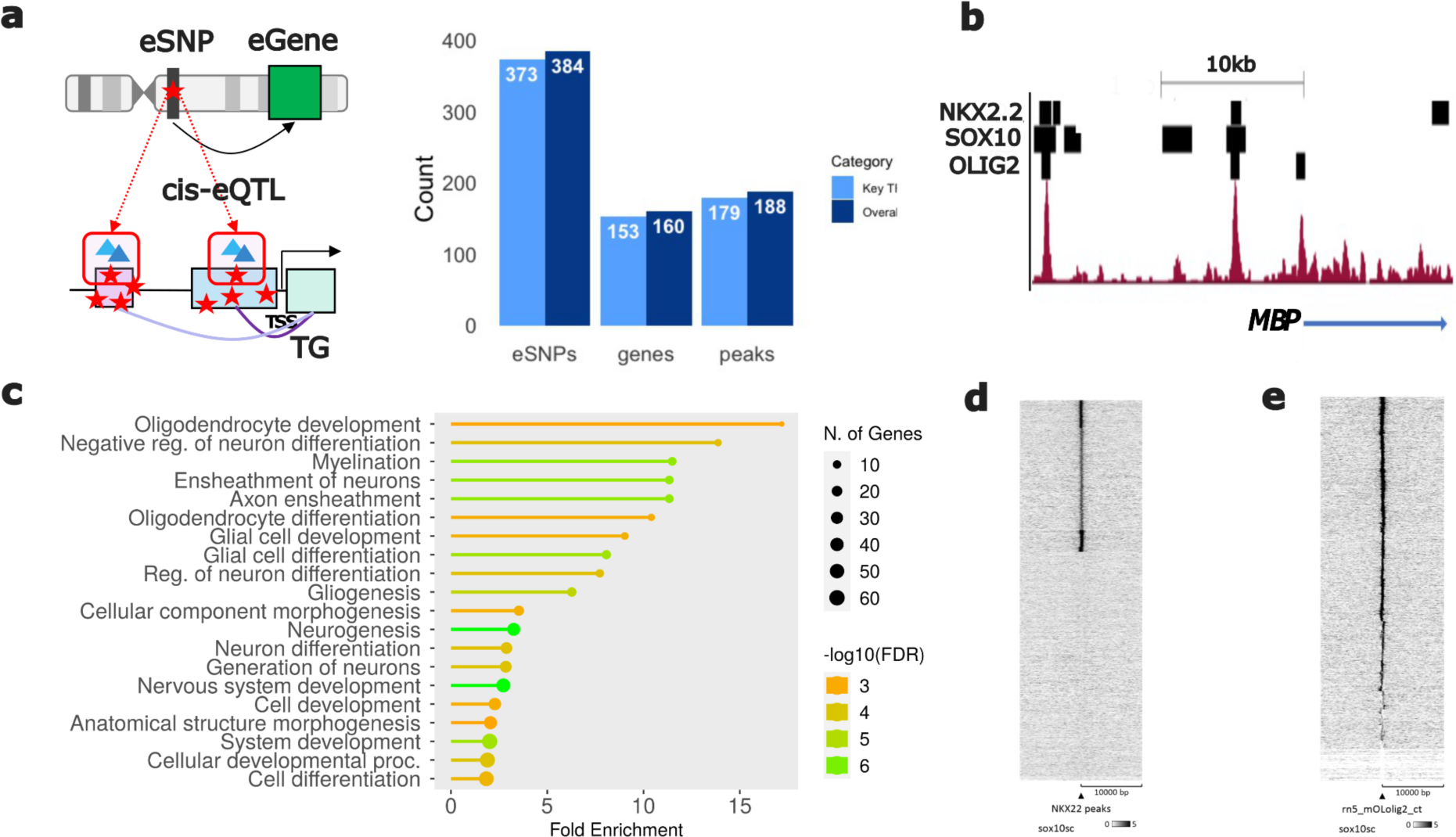
Independent validation using eQTLs and ChiP-seq data. **a** eQTL mapping onto oligodendrocytes regulatory regions **b** ChIP-seq peak for cooperative TF pairs that co-regulate *MBP*. The ChIPseq profile is for SOX10, and solid blocks indicate called peaks for the specified transcription factors. **c** Gene ontology analysis of target genes associated with oligodendrocyte eQTL’s, and **d,e** Heatmaps show distribution of SOX10 ChIP-seq reads centered on the previously defined NKX2.2 and OLIG2 sites.

### Validation of cooperative TF pairs

The model generated from human epigenome and expression data predicted a number of enriched TF pairs within oligodendrocyte-specific TF regulatory elements. In order to test if the coordination occurs as predicted, we utilized rat oligodendrocyte ChIP-seq data that were available for selected transcription factors. One predicted pair was OLIG2/SOX10, which had previously been shown to be extensively colocalized in analyses of rat oligodendrocytes^40^. To visualize the preferential binding of SOX10 on a global scale, a read density plot for SOX10 ChIP-seq reads^11^ was generated centered on the previously defined OLIG2 peaks^40^ in oligodendrocytes. In line with previous analysis, the average read density of SOX10 is highly enriched over OLIG2 bound sites. A novel pair predicted by the model was that of NKX2.2 and SOX10, and we generated a similar plot of SOX10 ChIP-seq reads over a defined set of NKX2.2 ChIP-seq peaks in oligodendrocytes^41^, and we found a similarly high enrichment of SOX10 binding on ∼40% of NKX2.2 binding sites. An example of the colocalization is shown for the *MBP* gene, which is highly expressed in oligodendrocytes. As shown, there are at least 2 sites upstream of MBP where there is colocalization of SOX10 with NKX2.2 and OLIG2.

## Discussion

With resources provided by advances in single-cell sequencing, some studies^42–45^ have elucidated the roles of several TFs, enabling the construction of cell type-specific gene regulatory networks to explain potential TF-TG relationships using bioinformatic tools. However, most of these studies and tools primarily focus on relationships between independent TFs and TGs.

This study identified co-binding TFs and their TGs in oligodendrocyte-specific regulatory regions. Deep learning models predicted TG expression levels using the expression levels of co-binding TF pairs, and we computed TF SI scores to define highly interacting co-binding TF pairs as ‘cooperative’ TFs that co-regulate TG expression levels. We found that the key co-binding TF pairs tend to highly interact with each other compared to non-key co-binding TF pairs for predicting TG expression levels. Independent validation, such as mapping eQTLs onto the regulatory regions, provides evidence for causal relationships between co-binding TF pairs and TGs. Additionally, converting these regions to the rat genome assembly coordinates and measuring the density of ChIP-seq signals for key cooperative TFs show that many of these TF pairs are enriched in the regulatory regions, indicating their collaborative role in co-regulating TG expression levels. We defined specific key TFs and examined co-binding TF pairs containing them, along with their interactions in predicting TG expression levels. We then compared these results with those of non-key TF pairs. Overall, co-binding TF pairs with known regulators of oligodendrocyte development exhibit higher SI scores, suggesting that they not only regulate TG expression individually but also cooperatively. We identified several highly cooperative TF pairs, such as SOX10 and OLIG2^12,46^, which are already known. Additionally, we discovered previously unreported cooperative pairs, such as SOX10 and NKX2.2, to the best of our knowledge.

Our study demonstrates several strengths. First, we concentrate on interactions between co-binding TF pairs and their impact on TG expression using deep learning approaches. Deep learning can elucidate complex TF relationships and their effects on TG expression levels. Second, our framework is straightforward and easy to apply. The pipeline’s code is openly available on GitHub, allowing users to input their scATAC-seq and scRNA-seq data for their specific purposes. Third, we provide a comprehensive analytical framework that incorporates analyses utilizing co-bindings by motif and expression levels. We define ‘cooperative’ TF pairs as TF pairs significantly co-enriched across regulatory regions, exhibiting high SI scores in terms of expression when predicting TG expression. The term cooperativity has often been applied to co-bindings of TFs to nearby sites that facilitates stabilized binding due to protein-protein interactions, but in our model, we use TF pairs that can bind to sites >50 bp apart, since TF’s can coordinately activate enhancers without direct interactions.

Nevertheless, there are some limitations to our study. To begin with, it’s important to note that more than two TFs can co-regulate TG expression^47,48^. However, our current tool is limited to analyzing interactions between two co-binding TFs. In future research, developing or applying more sophisticated methods capable of handling clusters of TFs that co-regulate the same TG expression will be informative. Moreover, our method for identifying binding sites relies on the position frequency matrices in the motif database. While both SOX10 and MYRF are key TFs for oligodendrocytes, we encountered difficulty in obtaining sufficient binding sites for MYRF. Consequently, we had to supplement with a different motif for MYRF based on our prior knowledge. More generally, the definition of TF motifs relies on disparate methods, and limitations of motif generation and analysis have been noted previously. Nonetheless, our analysis provided TF-TF coordination that we could validate using data from previous studies. We predict that future analysis can be used to determine if the predicted TF pairing plays a role in oligodendrocyte differentiation, since reliance on single factor studies is not able to recapitulate the important combinatorial functions of TF’s in generating cell type-specific gene expression patterns.

## Methods and Materials

### Pipeline for identifying cooperative TFs in oligodendrocyte gene regulation

First, we used published scATAC-seq data^21^ with peak-to-gene links. Second, we identified TFBSs and co-binding TF pairs from motif co-occurrence and co-enrichment analyses in oligodendrocyte regulatory regions. Third, we trained DNNs to predict TG expression levels and measure interaction effects between co-binding TFs using scRNA-seq data^22^, computing SI scores to identify cooperative TF pairs. Fourth, we built a gene regulatory network based on these interaction scores. Lastly, for independent validation, we mapped oligodendrocyte eQTLs onto regulatory regions with cooperative TF pairs, and performed co-enrichment analysis after converting these regions to the rat genome assembly to measure ChIP-seq signal density.

**<u>Step 1: Infer transcription factor binding sites</u>**

We inferred transcription factor binding sites (TFBSs) in 787 scATAC-seq peak regions that have linkages with TGs.

a. We used the R package *GenomicRanges* to format the ATAC-seq peaks into genomic ranges.
b. We set the position frequency matrices (PFMs) for the 949 motifs in *JASPAR2022* database^49^ in R and added nine more PFMS for the important modified motifs based on our prior knowledge.
c. We inferred TFBSs in the scATAC-seq peak regions using a R package, *motifmatchr*^50^.

**<u>Step 2: Identify co-binding transcription factor pairs</u>**

We identified co-binding TF pairs using the inferred TFBSs in Step 1.

a. We considered all possible TF-TF pairs that have binding sites in the scATAC-seq peak regions.
b. We removed TF pairs from the same TF families.
c. Co-occurrence analysis: We found TF pairs that have overlapping regions.
d. Co-enrichment analysis: We conducted hypergeometric tests to find significantly enriched TF pairs in the same regions. We used multiple testing corrections via false discovery rate (FDR) and applied FDR <0.1 cutoff. We define the TF pairs that are co-enriched (FDR < 0.1) as ‘co-binding’ TF pairs.
e. We built gene regulatory networks (GRNs) for TG-co-TF pair-peak links and matched TGs and co-TF pairs to the scRNA-seq data.
f. We removed lowly expressed TGs and TFs by applying a cutoff, median expression level > 1; more than half of the cells are expressed, from the GRNs.
g. We implemented differential expression testing using Seurat^51^ and selected TGs that are oligodendrocyte specific in the GRNs.
h. We annotated peaks, categorizing them into promoters and enhancers using *annotatr*^52^.

**<u>Step 3: Measure cooperativity of co-binding transcription factors</u>**

Gene expression levels of the co-binding TF pairs from scRNA-seq data were incorporated into deep learning models to predict the expression levels of the TGs and measure interaction effects between co-binding TFs on the expression levels of TGs using Shapley interaction (SI) scores.

a. We projected metacells for the cells in scRNA-seq data using a Python package, metacells^53^.
b. We inputted expression levels of TFs that have co-binding TFs and TGs to construct deep learning models for each TG using *PyTorch*^54^ in Python.
c. We computed SI scores for TF pairs in each deep learning model.
d. We created interaction matrices for the SI scores in deep learning models and calculated the mean interaction scores for co-binding TF pairs.
e. We calculated coefficients of variance (CV)^55^ of the interaction scores for each co-binding TF pair and removed the pairs that had CV values higher than 0.5.

**<u>Step 4: Gene regulatory network and TF hierarchy analysis</u>**

A gene regulatory network was built for six key cooperative TFs.

a. We chose one cooperative TF pair for each of the six key TFs based on the top interaction scores.
b. We built a gene regulatory network that has linkages for cooperative TF pairs to TGs.
c. We selected TGs that are co-regulated by cooperative TF pairs.
d. We generated a network plot using Cytoscape^56^.

**<u>Step 5: TF hierarchy analysis</u>**

TFs that can be TGs were chosen and we implemented hierarchy analysis^57^ for those TFs.

a. We calculated in-degree (I) and out-degree (O) for the TFs.
b. We computed a hierarchy height metrics for the TFs.
c. We defined TFs as top-regulator, middle-regulator, and bottom-regulator.

**<u>Step 6: Independent validation (eQTL, rat genome, and ChIP-seq)</u>**

We mapped the significant (FDR<0.05) oligodendrocyte eQTLs onto the scATAC-seq peak regions.

a. We downloaded publicly available oligodendrocyte eQTL data^39^ and subtracted the significant (FDR<0.05) eQTLs.
b. We mapped the significant eQTLs onto the scATAC-seq peak regions in our GRNs.
c. We verified our results by comparing the number of eQTLs mapped onto the peak regions for key-TF pairs and non key-TF pairs.

We performed the LiftOver analysis to convert genome coordinates for rat to human hg38 assembly using UCSC Genome Browser^58^.

a. We converted the genome coordinates for human hg38 assembly to the rn5 rat genome coordinates for human (hg19) assembly.
b. We found overlapping genome coordinates between conserved (from hg38 to rn5) assembly and the regulatory regions in our GRN.
c. We identified cooperative TF pairs in those overlapping regions found in b and the TGs they co-regulate.

Using the results from the LiftOver analysis, we tried to find signals in co-enriched binding sites for cooperative key TF pairs in rat oligodendrocyte ChIP-seq data. Heatmaps were created via EAseq^59^. ChIP-seq tracks were visualized using UCSC genome browser^58^. Previous ChIP-seq datasets for SOX10, OLIG2, and NKX2.2 are available at GEO accession numbers: GSE64703, GSE42447 and GSM1906296.

To verify the performance of deep learning model architectures, we trained a deep learning model for predicting a TG, *MBP* using another data^60^. We used the trained model to predict the expression level of *MBP* and compared the results with the model for *MBP* using the main data (**Fig S3**).

## Data and data processing

### Single-cell ATAC-seq

Chromatin accessibility data^21^ was used for the main analyses. Brain samples were selected and eight thousand nuclei from each sample were subjected to the Chromium Next GEM Single-Cell Multiome ATAC-seq. We filtered oligodendrocyte-specific peak-gene links for our analyses. 930 peaks and 606 genes were initially chosen.

### Single-cell RNA-seq

We downloaded scRNA-seq for the whole taxonomy collected from dorsolateral prefrontal cortex (1,395,601 cells) through the Open Data Registry on AWS as AnnData objects (h5ad format).^22^ We excluded cells for disease and kept the controls only. Then, we projected metacells for the whole taxonomy and found 2,004 metacells and 17,946 genes for oligodendrocytes.

### Uniform Manifold Approximation and Projection for Dimension Reduction

We gained scRNA-seq data for the whole taxonomy collected from dorsolateral prefrontal cortex through the Open Data Registry on AWS as AnnData objects (h5ad format)^51^. There were 1,395,601 cells across 18 sub-cell types. We selected 18,431 hg38 protein-coding genes, obtained using BioMart^61^, from among 36,517 genes. We normalized the data to a depth of 10,000 and log1 transformed it using Scanpy^62^ in Python. Then, we identified highly variable genes (HVGs) using dispersion-based methods^63^ to normalize dispersion, obtained by scaling with the mean and standard deviation of the dispersions for genes falling into a given bin for mean expression of genes. The cutoffs for the mean dispersions for genes were a minimum of 0.0125 and a maximum of 3, and for the minimum dispersion was 0.5. We identified 3,032 HVGs and scaled each gene to unit variance to clip values exceeding standard deviation of 10. To reduce the dimensionality of the data, we ran principal component analysis (PCA) and used top 30 PCs to compute the neighborhood graph of the cells. Finally, we embedded the neighborhood graph with 20 neighbors in two dimensions using Uniform Manifold Approximation and Projection for Dimension Reduction (UMAP)^64^.

### Differential expression testing

We inputted metacells for all cell types to identify oligodendrocyte-specific genes using Seurat v4 in R. We used the Poisson likelihood ratio test in FindMarkers function assuming that gene expression follows the negative binomial distribution. We grouped oligodendrocytes with oligodendrocyte precursor cells (OPCs) and astrocytes that are known as major cell types among glia in the central nervous system (CNS) and compared them to the other fifteen cell types. We used a cutoff, FDR (< 0.05) to select differentially expressed genes in the oligodendrocyte group.

### Position Weighted Matrices

Position weighted matrices (PWMs) for the 949 motifs in JASPAR2022 were used to infer transcription factor binding sites. We added PWMs for MYRF, SP7, and OLIG2 that are one of the key transcription factors from another study^65^, Mus musculus in JASPAR2022^49^, and HOCOMOCO v12^66^. We also included shorter motifs for other key TFs, such as SOX10, MYRF, ZNF24, NKX2.2, and SP7, considering their importance in oligodendrocytes (**Fig. S4**).

### Co-enrichment analysis

We used a hypergeometric test to assess whether a number of overlaps in the binding sites for two TFs follows a hypergeometric distribution. Specifically, given that a random variable *X* represents the possible outcomes of a hypergeometric process, the probability of getting k or more overlapping binding sites between two TFs inside a particular chosen set, as a hypergeometric random process, is

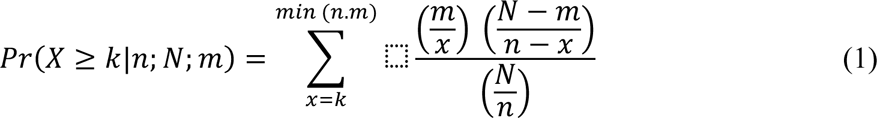

where *N* is the total number of transcription binding sites for all TFs, *m* is the number of binding sites for TF1, n is the number of binding sites for TF2, and *x* is the number of overlapping binding (co-occurrence) sites between TF1 and TF2. We applied an FDR adjusted p-value as a cutoff (< 0.1) for all possible TF pairs and chose co-binding TF pairs.

### Key transcription factors

We defined key TFs that are oligodendrocyte marker genes based on mouse loss-of-function studies that have shown that specific TF’s are critical for oligodendrocyte differentiation. This includes SOX10^23^, SOX2^24,25^, SOX8^26^, MYRF^27^, OLIG1^28^, OLIG2^29^, TCF7L2^30,31^, ZNF24^32^, NKX2.2^33^, and NKX6.2^34^ were chosen as key TFs.

### Deep learning models

We inputted expression levels of TFs that have co-binding pairs into the deep neural network (DNN) models to predict TG expression levels. 2,004 metacells (samples), 206 TFs (features), and a TG expression level (label) were used in the DNN models. We built a DNN for each TG to predict a TG expression level. The mean squared error (MSE) between predicted TG expression and actual TG expression was used as the loss function in DNN models. We cross-validated the training dataset (80% of the input samples) with 5-fold cross-validation and validated the best trained model on the 20% of hold-out validation dataset for the best use of data and to achieve reliable model performance. We used an early stopping function with patience 10 and determined the number of epochs and we set the batch size to 32. Adam with a learning rate 0.001 was used for training the models. The structure of our neural network model can be written as

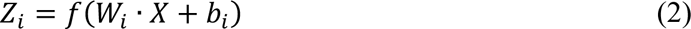

where *X* denotes the input data and *f* represents the activation function, specifically the LeakyReLU function. TF expression levels, *Z_i_* is the output of the *i*^th^ hidden layer, predicted TG expression level, and *W*_i_ and *b*_i_ are the weight matrix and bias vector for the *i*^th^ layer, respectively.

To evaluate the performance of our neural network model, we utilize the Mean Squared Error (MSE) loss function. The MSE quantifies the average squared difference between the predicted outputs of the model, Z and the true labels in our regression task. Mathematically, we can express the MSE as follows:

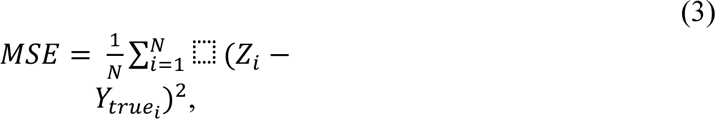

where *Z*_i_ represents the predicted output for the *i*^th^ sample, and *Y_truei_* denotes the true label corresponding to the *i*^th^ sample.

### Shapley interaction scores

We denote the set of all TFs by F, a feature i∈F, and a feature set S⊆F. We define the interaction effect between TF i and j, with feature set S, of a neural network f at a data point *X*_k_ to be

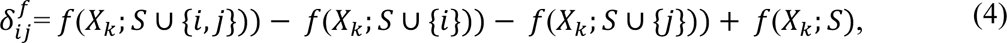

where *f*(*X*_k_; *S*) is the prediction at *X*_k_ when only TFs in S are used, which often requires retraining the NN multiple times. A common approximation is to replace the absent features (i.e., F\S) by the corresponding values in a baseline C_F\S_, such that

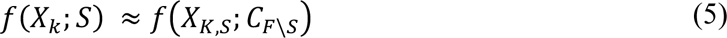

The baseline is set as the empirical mean of each feature. The Shapley interaction score 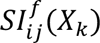 is the expectation of 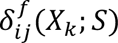,

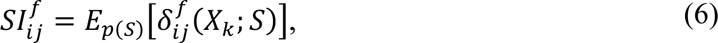

over a uniformly random chosen feature set *S* from *F*. We use Monte-Carlo procedure^67^ to approximate 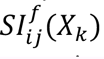 by a small number of samples of δ. To aggregate the local interaction effect at different data points into a global interaction effect, i.e., shared by the whole data domain, we use take the expectation 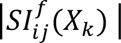 Q of w.r.t. the empirical data distribution *p*(*X*), such that

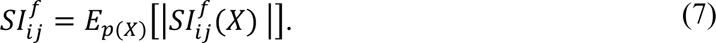

For a deep ensemble of our deep learning models, we used a posterior distribution of functions *q*(*f*) induced by the ensemble distribution of the weight *q*(*w*) where w is on Eq. (2). Thus, the interaction score is the expectation of 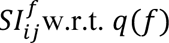, which we estimate by taking the average of *N*_3_ samples drawn from the ensemble:

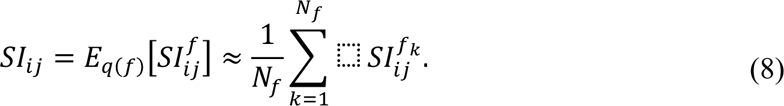

We computed Shapley interaction scores for the co-binding TF pairs, TF *i* and TF *j* using the trained DNN models and validation datasets. We calculated mean values for co-binding TF pairs using interaction matrices. We ranked them by percentile and scaled them to 0 and 1 for easier interpretation.

### Coefficient of variance

The coefficient of variation (CV) is a statistical measure of the dispersion of data points in a data series around the mean. The CV represents the ratio of the standard deviation to the mean, and it is a useful statistic for comparing the degree of variation from one data series to another, even if the means are drastically different from one another. The CV is defined as the ratio of standard deviation to the mean as follows:

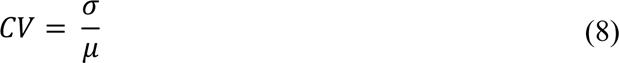

### Hierarchy analysis

We computed connectivity statistics, out-degree (O) and in-degree (I), for individual TFs to get a ‘hierarchy height’ metric (*h*), a normalized value of the difference between O and I for each TF. The ℎ is calculated as

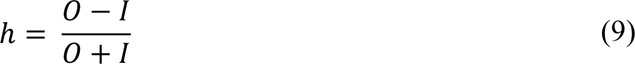

We defined TFs as top-regulator (h > 0.33), middle-regulator (−0.33 < h < 0.33), and bottom- regulator (h < -0.33) by their h values.

## Data availability

All data is publicly available on GitHub (https://github.com/jeromejchoi/scCo-reg).

## Code availability

The codes for the analyses and figures are available at https://github.com/jeromejchoi/scCo-reg.

## Acknowledgements

This work was supported by National Institutes of Health grants, R21 NS128761, RF1MH128695, R01AG067025, R21NS128761, and National Science Foundation Career Award 2144475, and a core grant to the Waisman Center from NICHD (P50 HD105353).

## Author information

### Authors and Affiliations

**Waisman Center, University of Wisconsin-Madison, Madison, WI, 53705, USA**

Jerome J. Choi, John Svaren & Daifeng Wang

**Department of Population Health Sciences, University of Wisconsin-Madison, Madison, WI, 53726, USA**

Jerome J. Choi

**Department of Comparative Biosciences, School of Veterinary Medicine, University of Wisconsin-Madison, Madison, WI, 53706, USA**

John Svaren

**Department of Biostatistics and Medical Informatics, University of Wisconsin-Madison, Madison, WI, 53076, USA**

Daifeng Wang

## Contributions

Conceptualization, J.S. and D.W.; Methodology, J.C., J.S., and D.W.; Formal Analysis, J.C.; Investigation, J.C., J.S., and D.W.; Writing – Original Draft, J.C.; Writing – Review & Editing; J.C., J.S., and D.W.; Supervision, J.S., and D.W.; Funding Acquisition, J.S. and D.W..

Correspondence to John Svaren or Daifeng Wang.

