## Supplementary Figures for "Single-cell multi-omics analysis reveals cooperative transcription factors for gene regulation in oligodendrocytes"

### **Supplementary information**

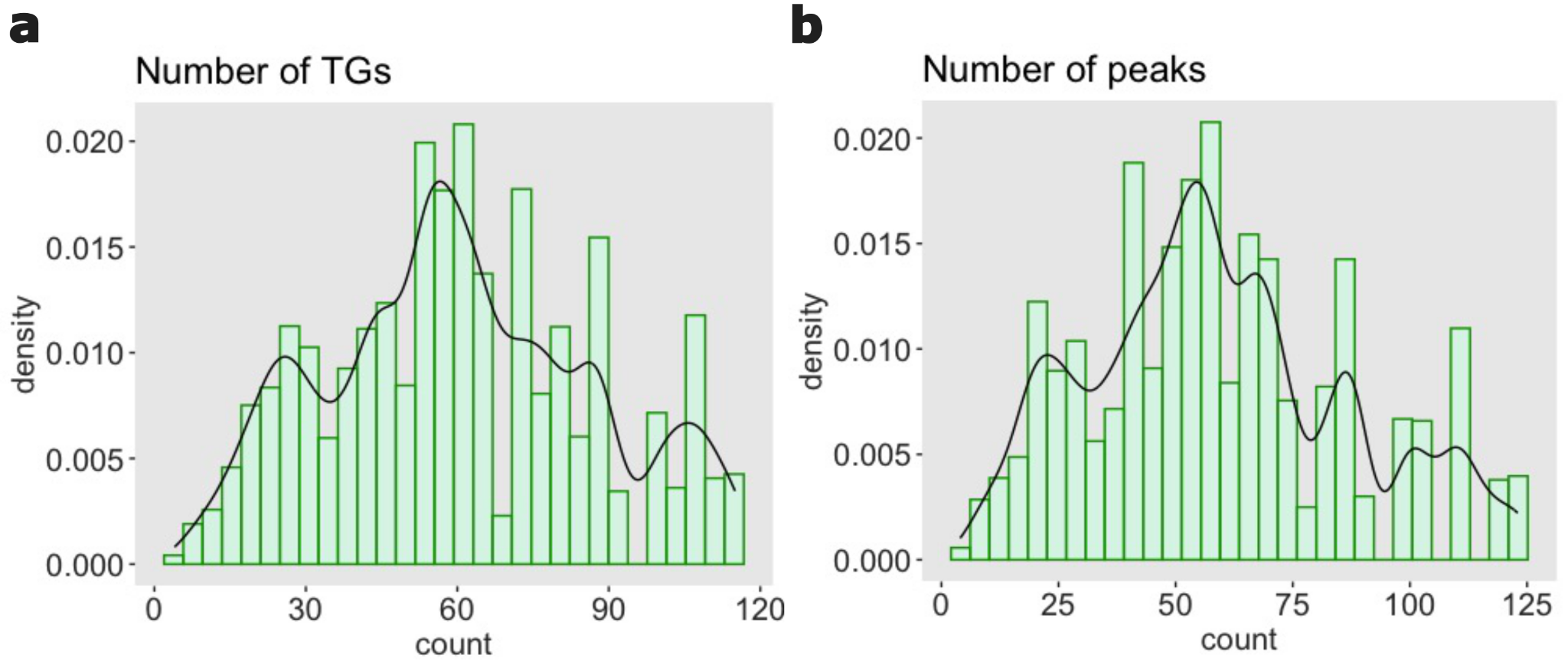

#### **Fig. S1: Distribution of the numbers of target genes and peaks for co-binding TF pairs. a** Distribution of the number of target genes (TGs) for co-binding transcription factor pairs. **b** Distribution of the number of peaks for co-binding transcription factor pairs.

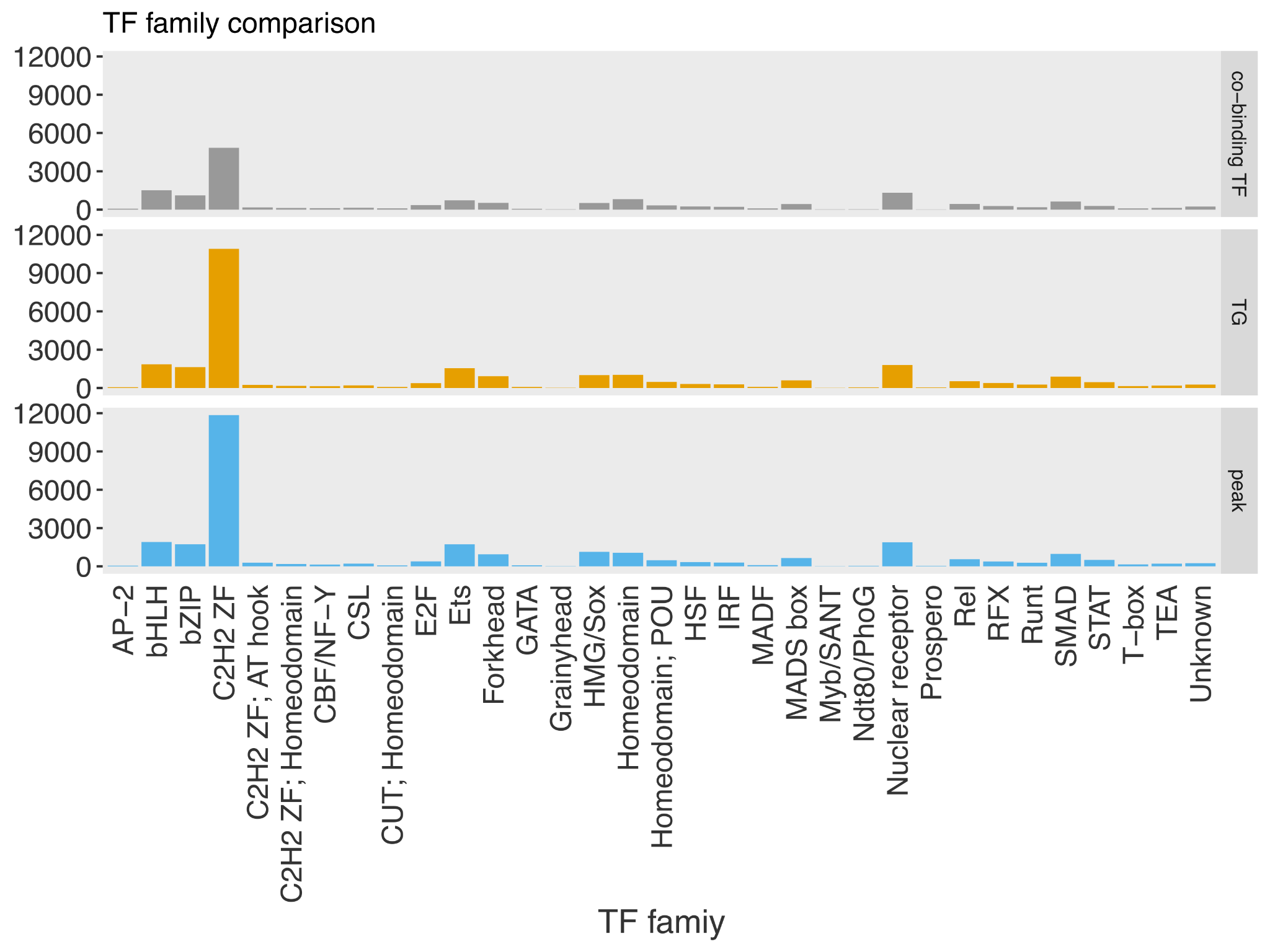

#### **Fig. S2: Transcription family comparison for transcription factors.** Distribution of the numbers of co-binding transcription factor (TF) pairs, target genes (TGs), and peaks for transcription factors by families.

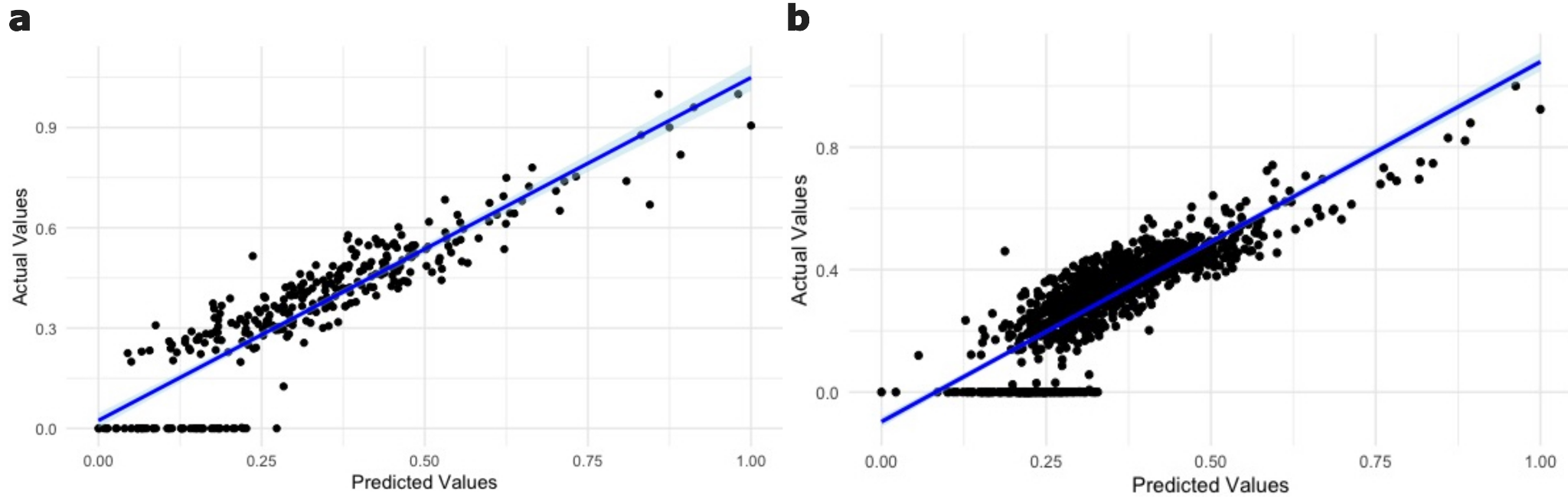

#### **Fig. S3: Validation for the deep learning prediction. a** Comparison between predicted values and actual labels in a deep learning model for predicting *MBP* using co-binding transcription factors in *Integrated Multimodal Cell Atlas of Alzheimer’s Disease* data. **b** Comparison *between* predicted values and actual labels in a deep learning model for predicting *MBP* using co-binding transcription factors in *Single-cell atlas reveals correlates of high cognitive function, dementia, and resilience to Alzheimer’s disease pathology* data.

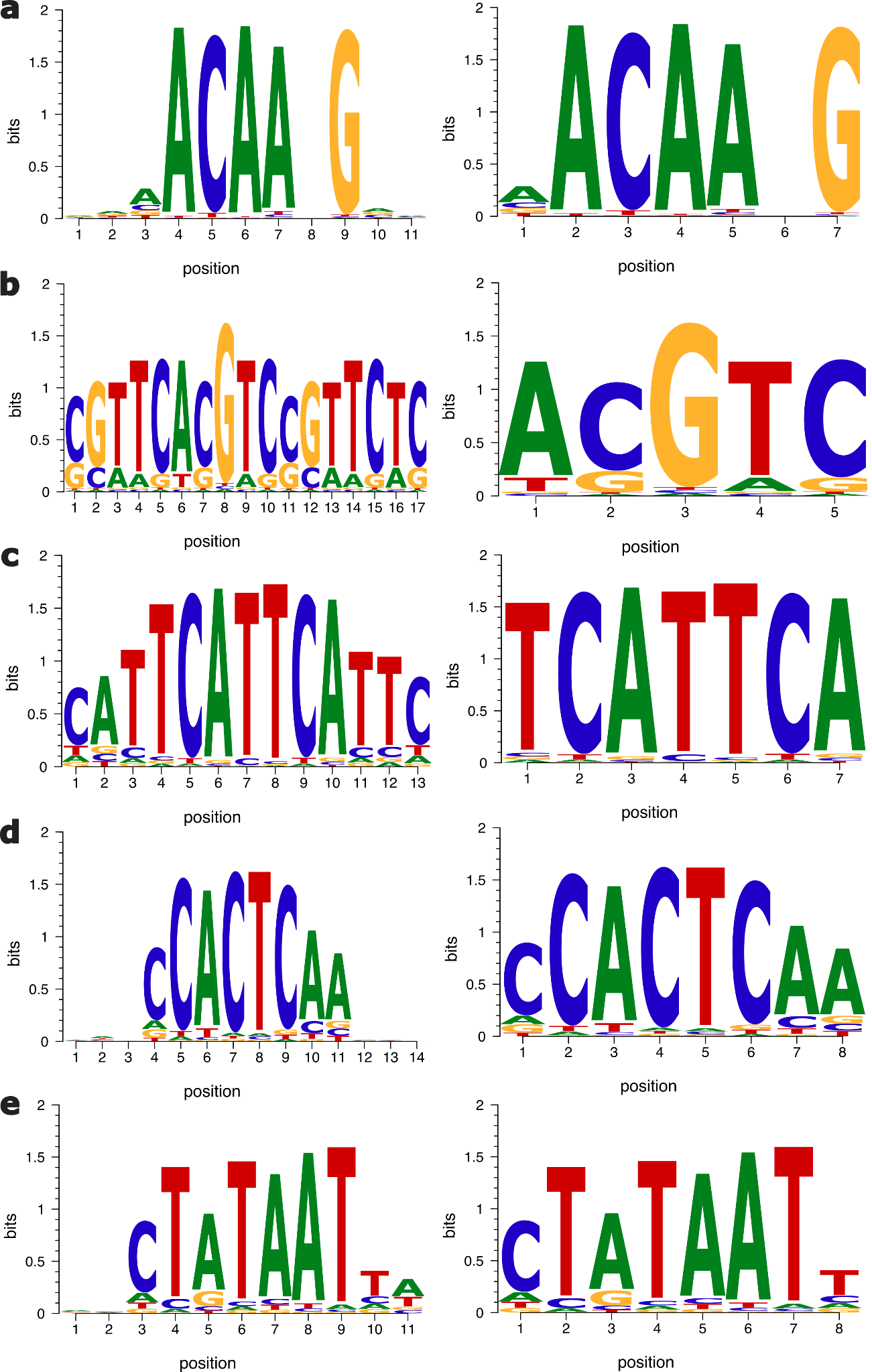

#### **Fig. S4: Original and shorter motifs. a** SOX10. **b** MYRF. **c** ZNF24. **d** NKX2.2. e SP7.

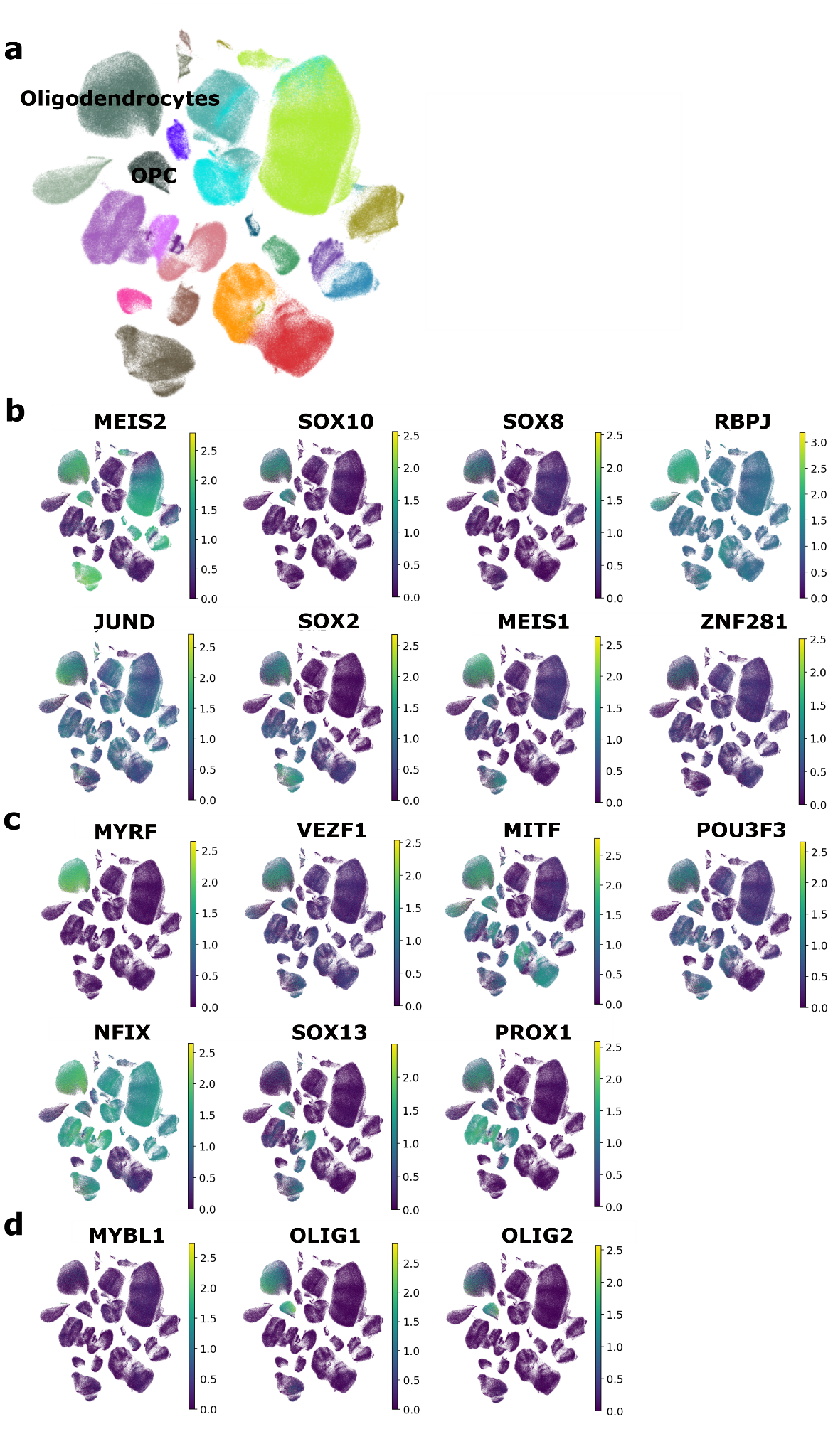

**Fig. S5: UMAPs for all cell types and TFs that can be TGs. a** top-level regulators (‘Master regulators’). **b** middle-level regulators. **c** bottom-level regulators.
